# Expression of the p53 Network Aging-associated Genes “shaken, not stirred” in a 3-Week Microgravity Simulation Human Study

**DOI:** 10.1101/2025.01.19.633763

**Authors:** Nik V. Kuznetsov

**Affiliations:** IBMP, UAEU

**Keywords:** aging, senescence, microgravity, p53 gene network, dry immersion, senolytics

## Abstract

**Background:** The hostile space environment affects human health and aging. NASA has named five main hazards astronauts will face in space, including space radiation and changes in gravity. However, the contribution of each of these factors, along with others, to the overall impact on biomolecular and cellular processes is not always clear.

**Objective:** To explore the effects of microgravity on the transcriptomes of healthy volunteers, with a focus on aging-related gene expression in p53 cell signaling pathways.

**Methods:** Ten healthy young men were exposed to simulated microgravity (SMG) for three weeks and blood samples were collected at five time point before, during and after the course of SMG. T cells were purified from the peripheral blood samples and total RNA was isolated and sequenced followed by bioinformatics analysis of volunteers’ global transcriptomes.

**Results:** A differential expression of p53 network genes was observed. The expression of nearly 30 genes involved in p53 gene network was affected during a 3-week course of simulated microgravity environment in dry immersion including classic p53 downstream target genes involved in cellular senescence: GADD45, p21, PUMA, IGF1 and other targets and potential target genes.

**Conclusion:** For the first time, the p53-associated cell signaling pathways and gene network in human T cells were reported to be affected *in vivo* by dry immersion SMG. It is evident that the relatively mild effects of simulated weightlessness on the human body are sufficient to activate these pathways and influence aging-related genes in the p53 gene network. These findings should not be dismissed, as they could open the door to the discovery of a novel category of drugs - MG-senolytics.

## 1. Introduction

The physiological changes caused by spaceflight are similar to those that occur with aging [1]. NASA’s Human Research Program has identified five hazards that astronauts will face in space. These include space radiation, isolation and confinement, distance from Earth, gravity (and the lack of it), and closed or hostile environments [2]. The use of a dry immersion (DI) microgravity simulation system is a well-established, ground-based approach to modelling of orbital space microgravity *in vivo*. It provides investigators with unique data on the physiological effects induced by microgravity alone [3]. The knowledge gained from those investigations is important for developing countermeasures against undesirable effects caused by space environment and aging [1]. The well-studied physiological impacts of microgravity on human organ systems still lack precise and thorough complements at the level of cellular effects and biomolecular response mechanisms to weightlessness alone.

The most commonly mutated gene in cancer [4], *TP53* tumor suppressor gene, which encodes the transcription factor p53, has been called the “guardian of the genome” by one of its discoverers, Professor Sir David Lane, [5] for its cell protective function under conditions of DNA damage stress.P53 gene network compiles multifunctional biomolecular players, including target genes downstream of p53. The p53 gene network responds to a variety of environmental and internal signals and comprises several crucial pathways including DNA repair, regulation of cell cycle ad induction of cell cycle arrest, apoptosis and cellular senescence. These pathways are vital for maintaining genome integrity and cellular homeostasis. p53 and several members of p53 gene network are considered to be involved in aging processes or mediating the longevity [6-8].

A four-fold increase was reported in the content of p53 in the skin cells of rats in space on STS-58 mission (Columbia) [9]. Furthermore, microgravity effects on the p53 pathway *in vitro* were registered in mouse sperm cells [10], cultured human lymphoblastoid cell lines [11,12], in macrophages [13] and in human soleus muscle after 3-day dry immersion [14]. Combinatory action of space radiation and orbital space microgravity during spaceflight led to increased mRNA levels of p53 in human lymphocytes of two healthy donors after 48 h on board the ISS [15].

## 2. Objective

In a dry immersion simulated microgravity study conducted on ten healthy volunteers, we aimed to investigate whether microgravity alone affects the expression of aging-related genes in the p53 gene network in human peripheral blood T cells *in vivo*.

## 3. Methods

### 3.1. Dry immersion study ethical approval

All participated volunteers signed the written consent according to the Helsinki Code of Medical Ethics for human samples. The Biomedical Ethics Committee of the IBMP RAS and Section of Physiology of the Bioethics Committee of the UNESCO National Bioethics Commission approved the study (Meeting number 483 took place on 03.08.2018).

### 3.2. Dry immersion experiment setup

An experiment with three weeks of exposure to simulated microgravity in dry immersion (DI) without any countermeasures was performed at the Institute of Biomedical Problems (Figure 1B) during a period of 8 months from September 2018 to April 2019 with the participation of ten healthy men aged from 24 to 32 years as described [16].

**Figure 1.**
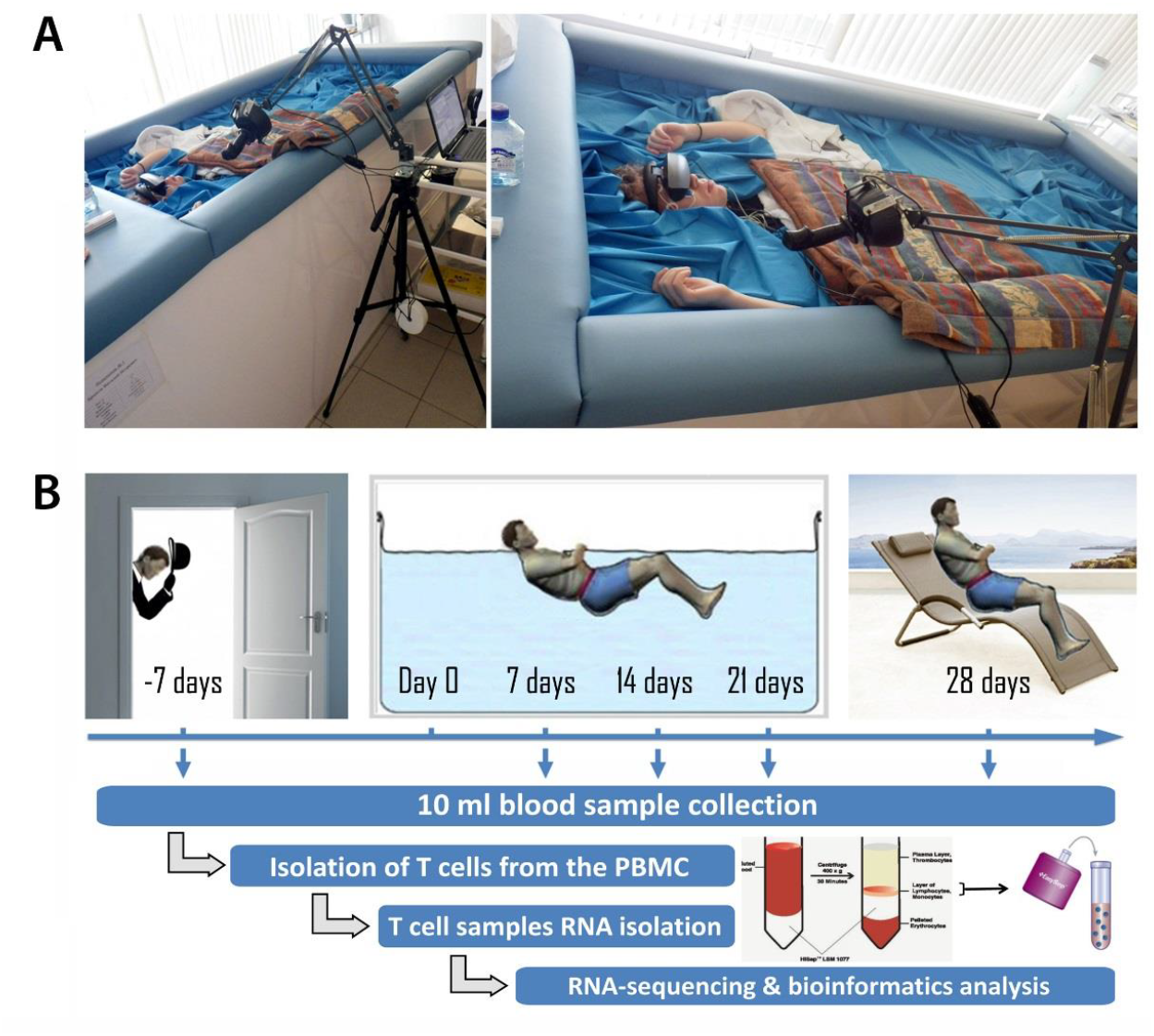
Setup of the simulated microgravity study in the dry immersion system. **A**. Dry immersion facility at the Institute of Biomedical Problems; **B**. The time course and flow chart of the simulated microgravity study.

### 3.3. Peripheral blood sample collection

In total, five samples of 10ml peripheral blood each were collected in sodium heparin tube from each volunteer at five time points during the time course of DI SMG study: at 7 days prior to Day 0 of DI (timepoint “-7d”; background, BG), after 7 days, 14 days and 21 days in DI SMG, and on the 7th day after DI SMG (timepoint “28 days”) (Figure 1B).

### 3.4. CD3+ T cell isolation

Peripheral blood mononuclear cells (PBMC) were isolated using Ficoll density gradient centrifugation isolation protocol optimised for Ficoll-Paque™ PLUS (Cytiva). Human T cells were enriched by the positive CD3 cell selection Kit (EasySep™ Human CD3 Positive Selection Kit II, Stem Cell Technology) with a purity of about 95%.

### 3.5. RNA Extraction

Total RNA was purified from sorted cells using RNeasy kit and treated with RNAse-free DNAse kit (##74104; 79254 QIAGEN, Hilden, Germany). The total RNA concentration was measured (NanoDrop2000, Thermo Scientific, Wilmington, DE, USA) and RNA integrity and purity was evaluated by gel electrophoresis analysis (1xTAE 1% Ultra Pure Agarose, #16500500 Invitrogen, Waltham, MA, USA). Gel images were created with ImageQuant LAS 4000 LAS4000 Image system using ImageQuant LAS 4000 Control Software (GE Healthcare, Freiburg, Germany) as described [17]. All selected isolated RNA samples were subjected to and have passed both the internal lab quality control (QC) and the QC performed by sequencing company (Novogene, Hongkong).

### 3.6. RNA-Seq and Bioinformatic Analysis

RNA sequencing and data quality control was performed with Illumina HiSeq-PE150 Platform at Novogene (Hongkong). Sequences were mapped to the reference genome with HISAT2, v.2.0.5. Reads were aligned to the human reference genome assembly December 2013 (GRCh38/hg38). Standard bioinformatics (filtering of data, alignment of RNA-seq reads, gene ontology) was arranged at Novogene (Hongkong). DESeq2 in R package was used for DEG analysis. In gene expression analysis, H-cluster, K-means, and SOM were used to cluster the log_2_ (ratios). Heatmap based on gene expression was composed in NASQAR (NYUAD). Functional classification of refined data for peak-related genes was performed using NCBI Gene resource www.ncbi.nlm.nih.gov (Bethesda, MD, USA) as described previously [18,19].

## 4. Results and Discussion

### 4.1. TP53 tumor suppressor gene expression

Transcript level of *TP53* tumor suppressor gene varied among all 10 volunteers in the study. At different timepoints of the experiment, *TP53* was up-regulated in 9 of 10 volunteers compared to background levels (timepoint “-7d”, Figure 1), and the dynamics of its expression varied between individuals. Seven volunteers (## 1-5, 7, 8) demonstrated an increase in *TP53* transcript level at 7 days in SMG and then showed variable expression with an overall upward trend. Volunteer #6 showed a slight decrease of *TP53* transcript level during the SMG course and then an up-regulation above background (BG) level at 28 days timepoint after SMG course. Volunteer #9 showed an overall down-regulation of *TP53* transcript with decline of the expression at 7 days and 14 days in SMG and then its level was partially recovered at 21 days timepoint. Volunteer #10 also showed a decrease in *TP53* expression at 7 days and 14 days in SMG, but it was increased on day 21 of SMG, and the expression level increased further after the SMG course on day 28 (Figure 2).

**Figure 2.**
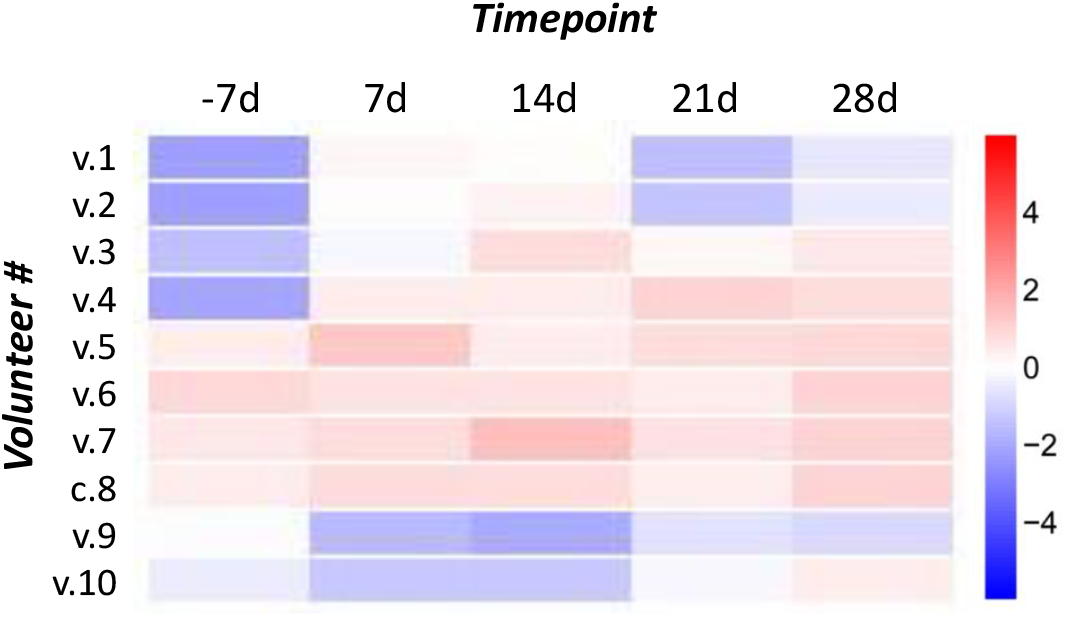
*TP53* gene expression in the dry immersion simulated microgravity study. *TP53* tumor suppressor gene is up-regulated in T cell samples of 9 out of 10 volunteers. Time course timepoints: before (−7d), during (7d; 14d; 21d) and after (28d) 3 weeks of simulated microgravity.

### 4.2. p53 gene network

The expression of nearly 30 genes involved in p53 gene network was affected during a 3-week course of simulated microgravity environment in dry immersion bed including classic p53 downstream target genes involved in cellular senescence and apoptosis: *GADD45, p21, PUMA, NOXA, p19ARF* and other p53 network associated and target genes. Genes that showed differential expression in all or nearly all of the ten volunteers in the study were listed (Table 1).

**Table 1.**
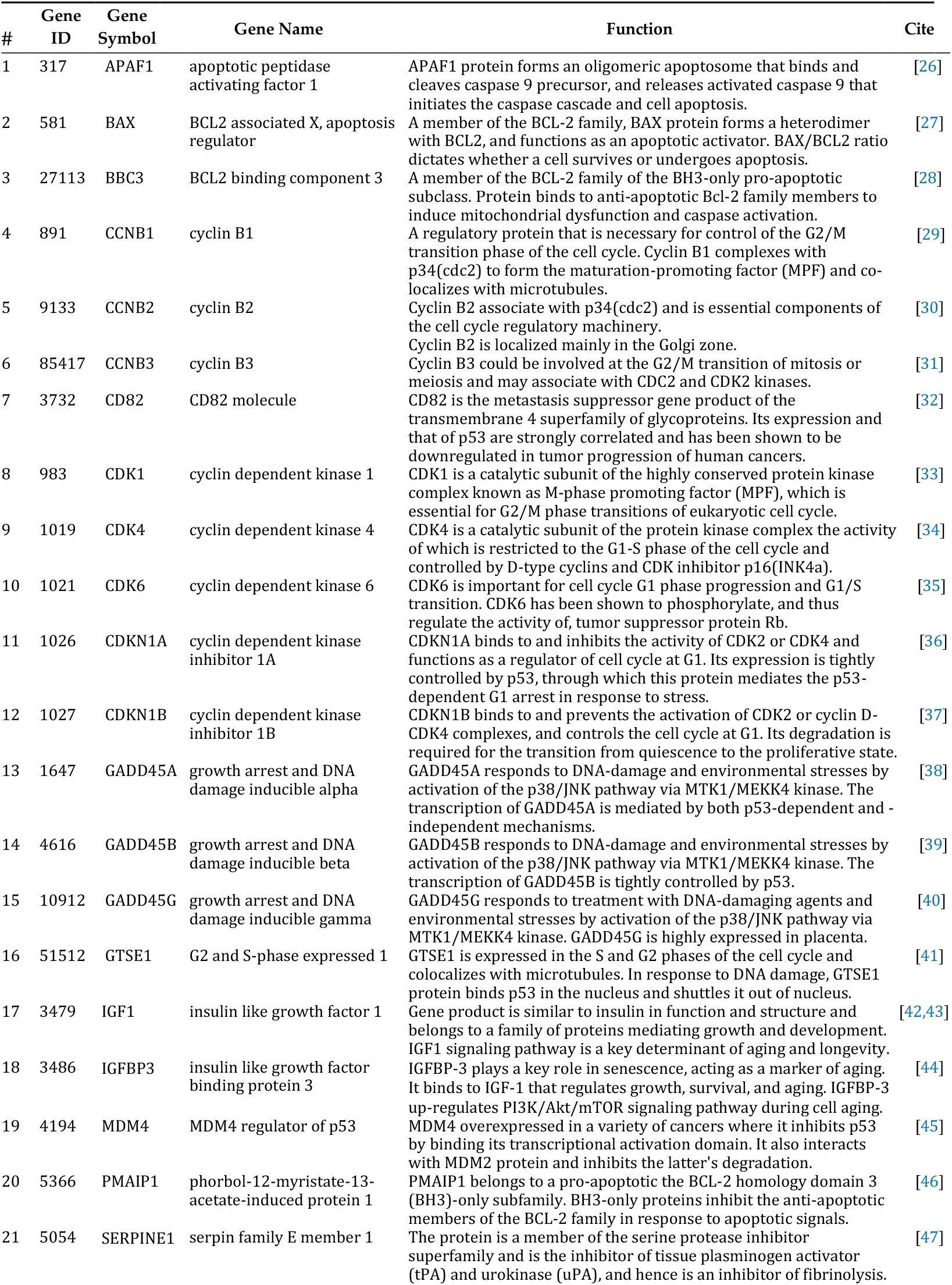
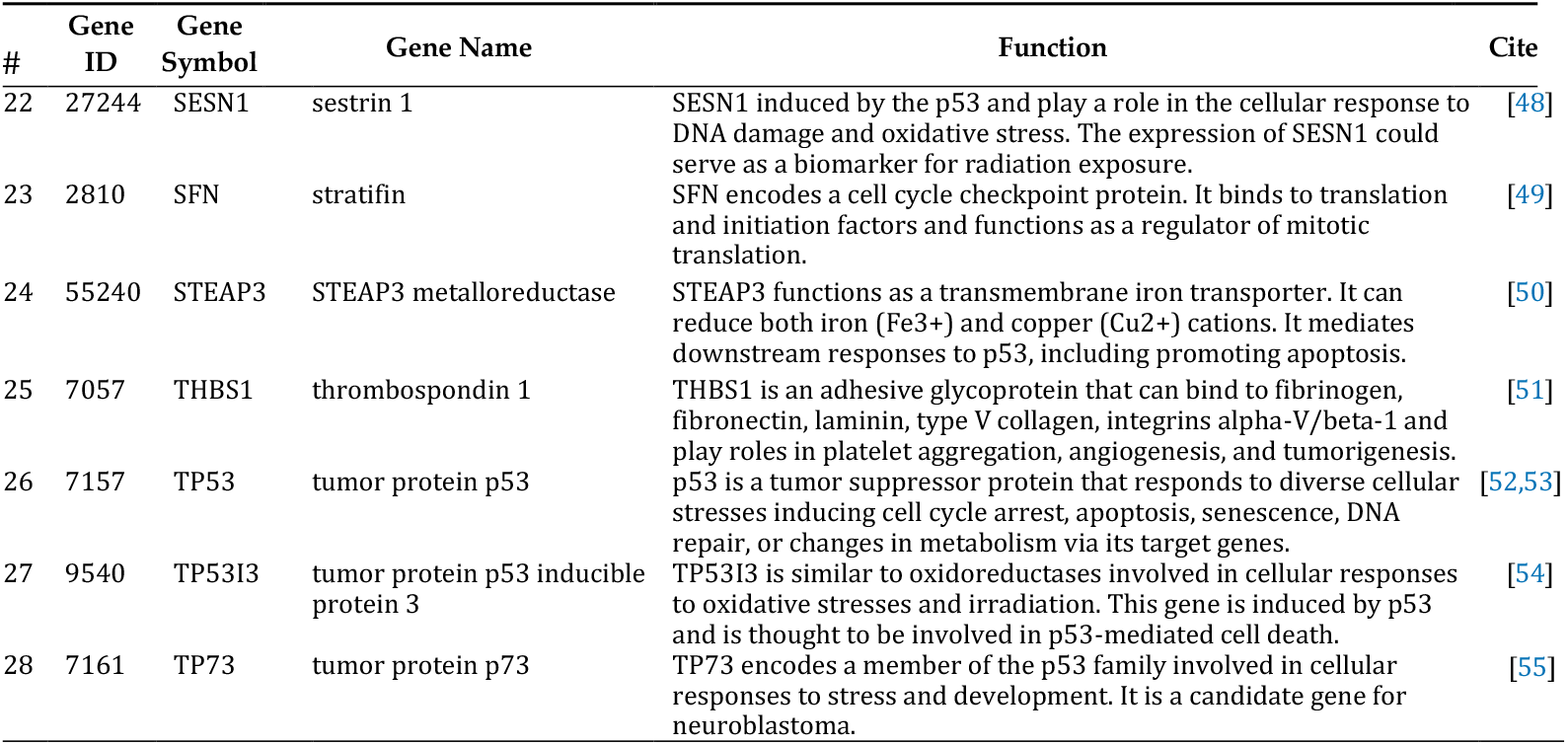
p53 pathway genes differentially expressed in 10 volunteers’ T cells during 3-week simulated microgravity study.

In relation to the regulation of the cell cycle, the gene expression of three B cyclins (CCNB1, CCNB2, CCNB3) and two cyclin dependent kinase inhibitors 1A and 1B in p53 gene network was found varying during DI SMG study in all ten volunteers.

Growth arrest and DNA damage inducible alpha, beta and gamma (GADD45A, GADD45B, GADD45G) playing roles in cell cycle arrest, DNA repair, apoptosis, innate immunity, genomic stability, and senescence showed coordinated differential expression during DI SMG study in all ten volunteers (Figure 3).

**Figure 3.**
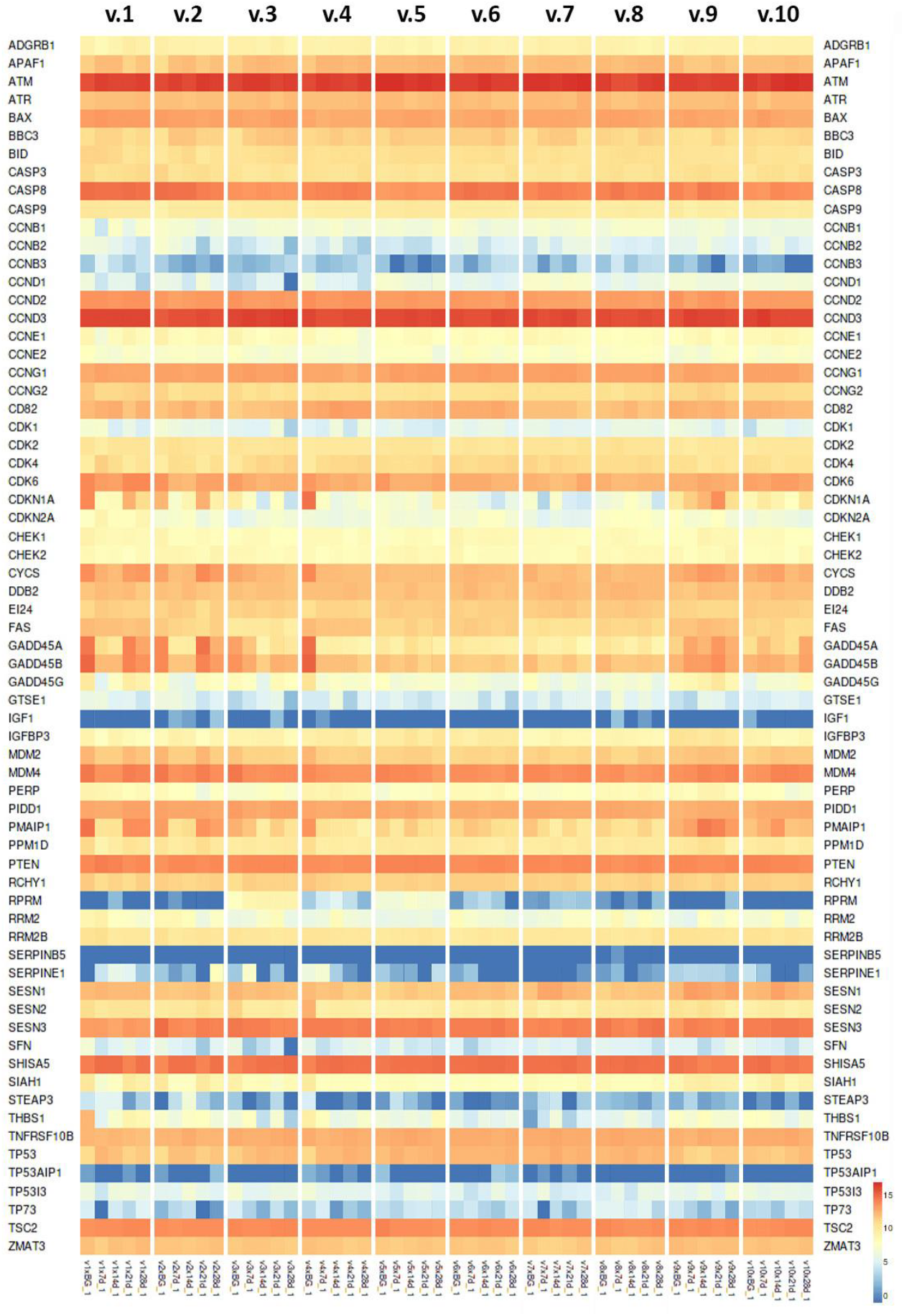
The heatmap of p53 gene network expression in DI SMG study. Expression of the genes involved in p53 gene network expressed in ten volunteers’ T cells at five timepoints: before (BG), during (7d; 14d; 21d) and after (28d) 3 weeks of simulated microgravity.

### 4.3. Aging-associated genes in p53 gene network

The differential expression of p53 network genes linked to aging and cellular senescence was recorded in the SMG study. The expression of dominant active p53 leads to constitutive expression of downstream target genes and results in premature aging [20]. There were varying changes in *TP53* transcript level observed during the experiment as described above (Section 3.1.).

The cell cycle cyclin dependent kinase inhibitor 1A (CDKN1A) interacts with cyclin dependent kinases CDK2 and CDK4 and blocks the progression through the cell cycle. BH domain Bcl-2-associated protein X (Bax) and p53 upregulated modulator of apoptosis (PUMA) are potent inducers of apoptosis. Cellular senescence and apoptosis prevent tumorigenesis. Also, both pathways have the potential to deplete stem and progenitor cell pools that leads to impaired tissue renewal ultimately impairing organ homeostasis – a hallmark of aging [21]. Cyclin dependent kinase inhibitors CDKN1A and CDKN1B, Bax, PUMA, cyclin dependent kinases CDK2 and, especially, CDK4 demonstrated the differential expression during DI SMG experiment in all ten volunteers (Figure 3). Sestrins are conserved stress inducible anti-aging genes [22]. The expression of three vertebrate sestrins was changed during DI SMG in all or almost all of ten volunteers.

In addition, p53 affects insulin-like growth factor 1 (IGF1) signaling pathway that is a key determinant of aging and longevity [21]. Reduced IGF-1 signaling is associated with extended lifespan in highly conserved pathway among different organisms from nematode to mammals [23]. IGFBP3 plays an important role in senescence as an aging marker. It binds to IGF-1 that regulates growth, survival, and aging. IGFBP3 up-regulates PI3K/Akt/mTOR signaling pathway during cell aging [24]. Also, p53 was elevated in IGFBP3 gene KO cells when compared to normal cells [25]. The gene expression of both IGF1 and IGFBP3 was noticeably changed during DI SMG study in 5 of 10 and in all 10 volunteers, respectively.

## 5. Conclusion

The principle of design, uniqueness and robustness of the dry immersion system allows for original research to be carried out in simulated microgravity conditions close to orbital ones. T cell response to simulated microgravity includes differential expression of p53 network genes. The involvement of aging-associated genes provides new insights into spaceflight practices and suggests the possibility of developing a new generation of microG-senolytics targeting p53 network genes.

## Author Contributions

N.K. contributed to the conceptual idea of the paper, formulated the objectives, wrote the manuscript, prepared the figures for data presentation and performed the editing.

## Acknowledgments

The author thanks colleagues at the New York University Abu Dhabi, Karolinska Institute, and Institute of Biomedical Problem RAS for the assistance and lively discussions. All interested participants of the project will be added as co-authors to the final version of the manuscript.

## Conflicts of Interest

The author declare no conflict of interest.

### Abbreviations

The following abbreviations are used in this manuscript:

BG: background
DI: dry immersion microG microgravity
PBMC: peripheral blood mononuclear cells QC quality control
SMG: simulated microgravity

